# Aldy 4: An efficient genotyper and star-allele caller for pharmacogenomics

**DOI:** 10.1101/2022.08.11.503701

**Authors:** Ananth Hari, Qinghui Zhou, Nina Gonzaludo, John Harting, Stuart A. Scott, S. Cenk Sahinalp, Ibrahim Numanagić

**Affiliations:** Department of Electrical and Computer Engineering, University of Maryland, College Park, MD, U.S.A.; Department of Computer Science, University of Victoria, Victoria, B.C., Canada; Pacific Biosciences, Menlo Park, CA, U.S.A.; Department of Pathology, Stanford University, Palo Alto, CA, U.S.A.; Cancer Data Science Laboratory, National Cancer Institute, National Institutes of Health, Bethesda, MD, U.S.A.

## Abstract

High-throughput sequencing provides sufficient means for determining genotypes of clinically important pharmacogenes that can be used to tailor medical decisions to individual patients. However, pharmacogene genotyping, also known as star-allele calling, is a challenging problem that requires accurate copy number calling, structural variation discovery, variant calling and phasing within each pharmacogene copy present in the sample.

Here we introduce Aldy 4, a fast and efficient tool for genotyping pharmacogenes that utilizes combinatorial optimization for accurate star-allele calling across different sequencing technologies. Aldy 4 adds support for long reads and ships with a novel phasing model and improved copy number and variant calling models.

We compare Aldy 4 against the current state-of-the-art star-allele callers on a large and diverse set of samples and genes sequenced by various sequencing technologies, such as whole-genome and targeted Illumina sequencing, barcoded 10X Genomics and PacBio HiFi. We show that Aldy 4 is the most accurate star-allele caller with near-perfect accuracy in all evaluated contexts. We hope that Aldy remains an invaluable tool in the clinical toolbox even with the advent of long-read sequencing technologies.

**Availability:** Aldy 4 is available at https://github.com/0xTCG/aldy.

## 1 Introduction

The rapid development of high-throughput sequencing (HTS) technologies has ushered in the era of precision medicine that aims to tailor medical decisions at the individual level (Hamburg and Collins, 2010). A key component of precision medicine is pharmacogenomics which studies the associations between the individual genotypes of clinically important genes (also known as *pharmacogenes)* and individual variation in drug response (Weinshilboum and Wang, 2017). While the HTS data theoretically provides sufficient means to accurately genotype any gene in a given individual, genotyping of many pharmacogenes remains practically challenging (Twesigomwe *et al*., 2020). One of the key challenges is the fact that many pharmacogenes of vital clinical importance—most notably the *CYP2D6* gene, whose genotype impacts up to 25% of clinically prescribed drugs (Ingelman-Sundberg, 2004)—are highly polymorphic, and furthermore located next to the highly similar pseudogenes due to being located within segmental duplications (Ingelman-Sundberg, 2004). These pharmacogenes are also subject to various copy number and structural changes, e.g. through a fusion event between a gene and its pseudogene, possibly due to the instability of the segmental duplication region wherein they reside (Sezutsu *et al*., 2013). These issues need to be carefully and comprehensively accounted for before the genotyping process, in order to obtain accurate results. Lastly, alleles of many pharmacogenes are not defined through a single nucleotide variant (SNV), but through a complete gene haplotype. Thus the exact functional impact of an allele can only be determined through phasing, or haplotyping, of the whole genic region. In the pharmacogenomics community, haplotyping is commonly known as *star-allele calling* (Robarge *et al*., 2007), owing to the fact that most of the known pharmacogenetic haplotypes are assigned a unique star-allele identifier.

Standard tools for genotyping HTS datasets, such as Genome Analysis Toolkit (GATK) (McKenna *et al*., 2010; Poplin et al., 2017), cannot be used for star-allele calling because they are unable to haplotype the whole genic regions and assign correct star-alleles. General-purpose computational phasing tools, such as HapCUT2 (Edge *et al*., 2017) and HapTree-X (Berger *et al*., 2020), are also inadequate for calling star-alleles: these tools are either designed for phasing diploid organisms and thus cannot phase regions that underwent significant copy number changes, or cannot handle fusions and other structural variations. Furthermore, the distance between allele-defining variants is often too large and, as a result, many alleles cannot be phased with short-read sequencing data. On the other hand, statistical phasing tools, such as BEAGLE (Browning *et al*., 2021) or Eagle (Loh *et al*., 2016), also do not handle the presence of fusions and structural variations. Thus most of the star-allele calling was— and still is— being done by various custom primer-specific PCR and array assays (Numanagić *et al*., 2015), mostly due to their price and speed, despite calls generated by these assays being often limited in breadth and scope (Pratt *et al*., 2010; Fang *et al*., 2014).

A number tools have been recently developed to address the challenge of accurate star-allele calling (Caspar *et al*., 2020). Cypiripi (Numanagić *et al*., 2015), the first tool specifically designed for this purpose, supported calling *CYP2D6* star-alleles from Illumina WGS data. Cypiripi was followed by Aldy (Numanagić *et al*., 2018), Stargazer (Lee *et al*., 2019), Astrolabe (Twist *et al*., 2016), StellarPGx (Twesigomwe *et al*., 2021) and Cyrius (Chen *et al*., 2021). These tools aggregate the data from the existing star-allele databases, such as PharmVar (Gaedigk *et al*., 2018), and use it to call star-alleles directly from HTS data. Some of these tools, such as Aldy and Stargazer, are also able to detect copy number changes and fusions with a high level of accuracy. However, the majority of these tools target only a small set of pharmacogenes (typically *CYP2D6* and other cytochrome P450 genes) and are tuned for short-read HTS data generated by the Illumina whole-genome sequencing (WGS), whole-exome sequencing (WES) and (in some cases) targeted sequencing panels such as PGRNseq (Gordon *et al*., 2016).

In recent years, we have witnessed a slow but steady shift toward third-generation HTS technologies such as PacBio and Oxford Nanopore (De Coster *et al*., 2021). These technologies produce significantly longer reads (typically measured in tens of kilobases) than Illumina reads (measured in tens of basepairs). While they were initially dismissed in clinical settings due to the high cost of sequencing and high error rates, these technologies are making a resurgence thanks to the recent improvements in terms of accuracy and cost. For example, PacBio HiFi sequencing offers up to 25 kbp-long reads with a 99.5% accuracy rate (Hon *et al*., 2020). Unfortunately, not many tools are able to correctly use the data generated by these technologies for calling pharmacogenomic star-alleles due to the different assumptions and biases as compared to the standard Illumina short-read data. Star-allele callers are also unable to make use of the long-range information within long reads for better phasing of allele-defining variants.

Here we present Aldy 4, the next iteration of Aldy software that addresses the aforementioned challenges. Aldy 4 completely revamps its original star-allele calling pipeline and adds support for long-read technologies such as PacBio HiFi. The changes include an alignment correction module that addresses various biases and errors common during the alignment of long reads to the pharmacogenomic regions. It also provides a novel star-allele calling pipeline that incorporates the long-range phasing information from long reads into the star-allele calling model. Finally, Aldy 4 brings support for 20 new pharmacogenes, provides an easy interface for adding the support for other pharmacogenes, adds an application programming interface (API) for easy incorporation of pharmacogenomic calling within the existing pipelines, and brings various other improvements to the original pipeline. We compare Aldy 4 against other popular star-allele callers such as Cyrius (Chen *et al*., 2021), StellarPGx (Twesigomwe *et al*., 2021), Stargazer (Lee *et al*., 2019), and Astrolabe (Twist *et al*., 2016) and Aldy 3 (Numanagić *et al*., 2018) on 20 pharmacogenes and four sequencing technologies (Illumina WGS, 10X Genomics, PacBio HiFi, and PGRNseq v.3) and show that its accuracy is better than or equal to that of the other tools. We also demonstrate that Aldy has minimal impact on system resources, typically needing only afew minutes to genotype and phase a sample without requiring the expensive pre-processing steps such as variant calling. We hope that Aldy will remain a crucial tool in the pharmacogenomics toolbox even with the advent of long-read sequencing technologies.

## 2 Methods

The goal of Aldy is to reconstruct the exact sequence content (or haplotype) of each gene copy of a given pharmacogene from a high-throughput sequencing (HTS) data sample and assign a star-allele to each reconstructed haplotype present in the dataset. This process is subsequently referred to as *star-allele calling*.

In order to accurately call star-alleles, it is necessary to consult a database of known star-alleles that contains the exact sequence content of each pharmacogene allele. Suppose that a pharmacogene *G* harbors variants 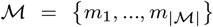, where any 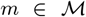 is a single nucleotide polymorphism (SNP) or a small indel. Depending on their impact on the gene *G*, these variants are either functional or silent. The reference allele of *G*, typically known as *1 star-allele, is an allele that harbors no variants at all. Any other star-allele *S_i_* is defined by the subset of known variants 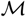 that distinguish its sequence content from the reference *1 allele.

In some genes, such as *CYP2D6*, star-allele identifiers are also assigned to fusions and other pseudogene-induced structural changes that affect the pharmacogene. For this reason, we need to extend the definition of star-alleles to also include their structural configuration. This configuration describes whether a pharmacogene is wholly present in the genome, deleted, or is a gene-pseudogene hybrid. The set of valid configurations is denoted as 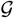. Note that each structural configuration can induce many distinct star-alleles depending on the choice of mutations from 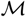. Thus, we can formally define a *star-allele S_i_* as a tuple (g_*i*_, *A_i_*) where 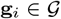 and 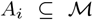. The *star-allele database* is formally a collection of all known structural configurations, mutations and known star-alleles 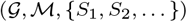 where *S_i_* = (g_*i*_,*A_i_*) such that 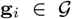 and 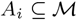.

To call star-alleles of a given pharmacogene from the given sequencing sample, Aldy needs to perform the following steps:

- analyze the aligned HTS reads in BAM/CRAM format and resolve incorrectly aligned reads;
- detect structural configurations by calling copy number changes and gene-pseudogene fusions; and
- use the read alignments from BAM/CRAM to call star-alleles and phase the gene.

### 2.1 BAM/CRAM analysis

Aldy begins by taking a SAM, BAM or CRAM file (Li *et al*., 2009) generated by a read aligner (e.g. `BWA` (Li and Durbin, 2009), `pbbm2`^1^ or `minimap2` (Li, 2018)). It is recommended to post-process these files with the GATK’s “Best Practices” pipeline (Van der Auwera *et al*., 2013) (the local indel realignment step is especially helpful for the subsequent variant calling). Aldy extracts the relevant variants that are present in a given pharmacogene from the alignment file, as well as coverage information needed for the copy number and structural variation detection step. It also collects phasing information from long reads, barcoded fragments and paired-end fragments where available.

The original version of Aldy relied on the assumption that read alignments produced by the off-the-shelf aligner are mostly correct. While this assumption holds for short paired-end Illumina reads, it breaks for long reads such as PacBio HiFi reads. For example, if a sample harbors a gene duplication and if the highly similar pseudogene is located immediately next to this gene, any long read spanning two duplicated copies of the gene will get its second half incorrectly aligned to the pseudogene because the reference genome does not contain two copies of the gene in question (Figure 1). The correct alignment would perform a split mapping and align the second half again to the pharmacogene. These incorrect alignments are even more problematic in the presence of gene fusions: any read that spans a gene-pseudogene fusion breakpoint will not be split-mapped but incorrectly aligned to either pharmacogene or its pseudogene.

**Fig. 1.**
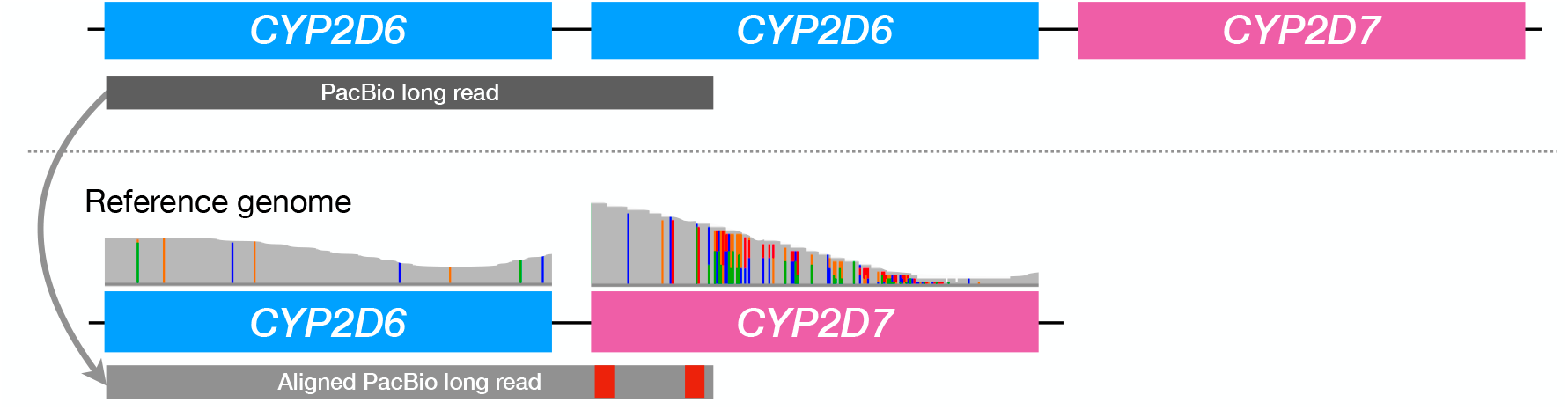
An example of an incorrect long read alignment to the reference genome. If a donor genome (above) contains two copies of CYP2D6 pharmacogene, any long read (gray rectangle) that spans both copies will get aligned to the reference genome (below) that contains only a single CYP2D6 copy. However, this read will get its second half (containing CYP2D6 sequence) incorrectly aligned to the CYP2D7 pseudogene due to the high sequence similarity between these genes. The final result is the over-abundance of coverage in the pseudogene region as compared to the CYP2D6 region (an IGV coverage plot is shown above the reference genome).

Aldy 4 resolves this problem by splitting any long read that spans the gene-pseudogene boundary into shorter gene-level chunks and aligning each chunk independently. Each chunk is guaranteed to span only one gene (either pharmacogene or a pseudogene) and thus avoid being misaligned in the manner described above. Aldy 4 performs a further split-mapping of each chunk that spans a potential fusion breakpoint to determine whether a read originates from a fusion event or not (a read is said to originate from a fusion event if its split-mapping alignment score is lower than the original alignment score).

Unlike previous versions, Aldy 4 considers base quality scores and read mapping qualities when calling the allele-specific variants. This ensures that the low-quality variants in noisy and low-coverage samples are filtered out before the star-allele calling.

### 2.2 Copy number and structural variation analysis

In a typical scenario, a sample contains only two parental copies of a pharmacogene of interest for which star-alleles need to be called. This is true for most of the pharmacogenes of interest. However, a few major pharmacogenes do not follow this pattern and are prone to various copy number changes and structural events. The most notable example is that of the *CYP2D6* gene, perhaps the most important of all pharmacogenes (Ingelman-Sundberg, 2004), whose copies can undergo whole gene deletions, duplications and hybrid fusions (where a copy begins with the *CYP2D6* sequence but switches to the pseudogene *CYP2D7* sequence at a given breakpoint, or vice versa) (Kramer *et al*., 2009). Each copy—fusions included—yields its own star-allele. Thus, in order to correctly call star-alleles of such genes, it is necessary to correctly detect the total number of available gene copies as well as the configuration (i.e., structure) of each copy.

Each gene copy can be described by its structural configuration represented as a binary vector 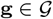 that indicates the presence or absence of genic regions in a given configuration (Figure 2). Because each star-alleleis defined by a matching structural configuration, such configurations must be found before the star-alleles can be accurately called. The size of the configuration vector depends on the number of gene segments that define various structural configurations. For example, the *CYP2D6* gene is divided into *r* = 20 segments that correspond to its exons, introns and flanking regions, because all structural variations are described at the level of whole exons and introns (Kramer *et al*., 2009). The total length of the *CYP2D6* configuration vector is 2*r* (i.e. 40) because the vector also includes segments from the neighboring *CYP2D7* pseudogene. This vector can encode any known *CYP2D6* structural configuration: for example, a single *CYP2D6* copy (*r* ones followed by *r* zeros), a single *CYP2D7* copy (*r* zeros followed by r ones), *CYP2D6–2D7* fusion in intron 1 (one followed by *r* – 1 zeros, in turn followed by a 0 and *r* – 1 ones) and so on. Once these vectors are established, any complex configuration within CYP2D locus can be represented as an aggregate of individual configuration vectors (see Figure 2 for details).

**Fig. 2.**
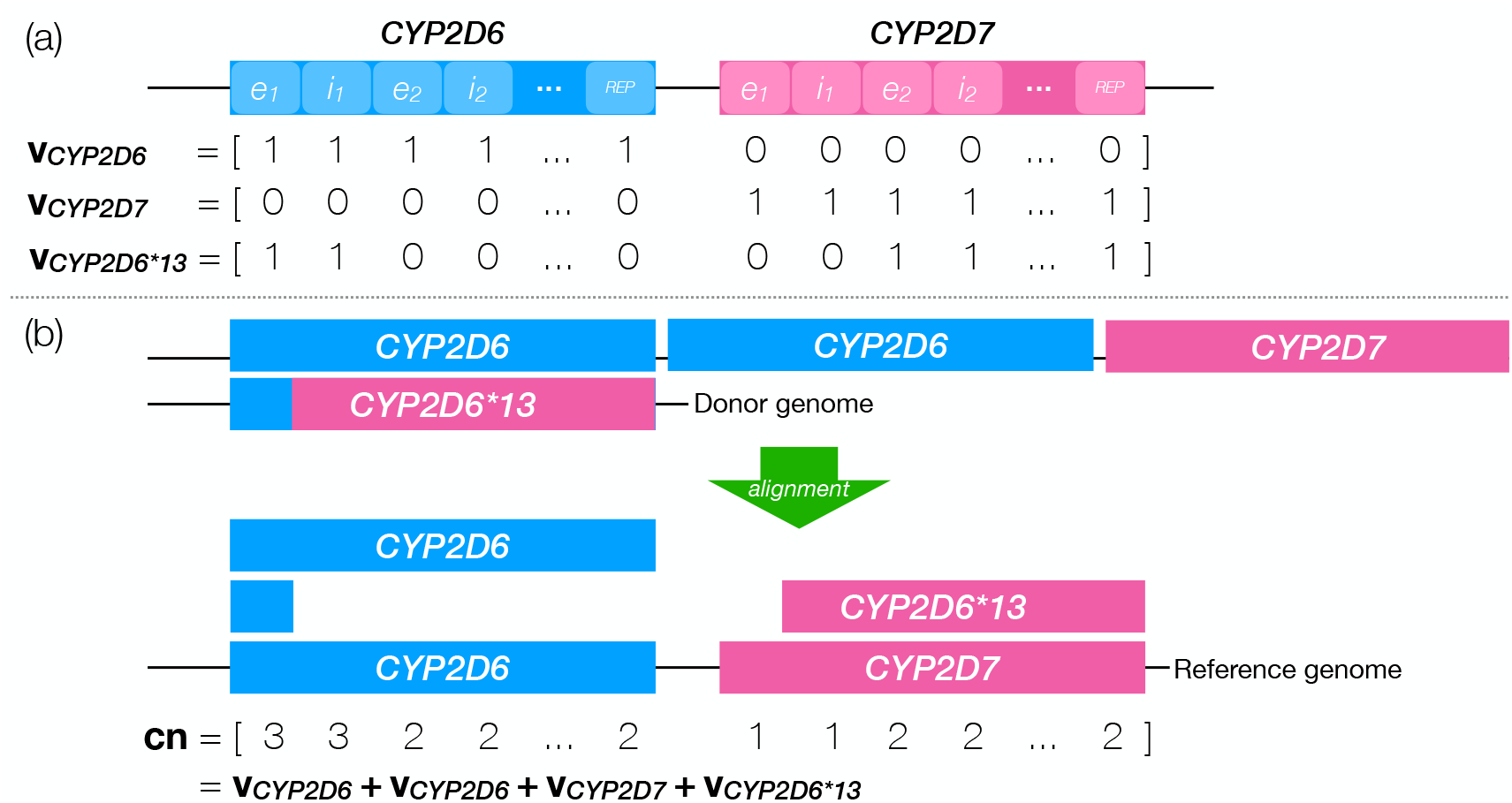
(a) An example database of CYP2D6 structural configurations containing three such configurations (**v**_CYP2D6_, v_CYP2D7_ and **v**_CYP2D6*13_). Regions on top of which the configurations were defined (i.e., *e*_1_, *i*_1_ etc.) are shaded with lighter color. (b) Sample decomposition of the aggregate coverage vector **cn**, observed after aligning the reads originating from the donor genome (above) to the reference genome (below). As can be seen, **cn** can be expressed as the sum of 4 structural configuration vectors from the database.

In a sequenced sample, we only observe the aggregate coverage vector **cn** that describes the number of reads covering each genomic loci of interest within the sample. The goal of Aldy is to find a set of configuration vectors 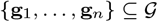 whose sumis closest to the observed aggregate coverage.^2^ As there might be many such sets, Aldy only looks for the most parsimonious solution: a solution that selects the minimal number of such vectors. This problem, previously dubbed as the Copy Number Estimation Problem (CNEP) (Numanagić *et al*., 2018), can be efficiently solved via integer linear programming (ILP) as follows.

Assume that a gene *G* is segmented into 2*r* regions. Let 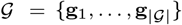 stand for the set of the available configuration vectors, where g_*i*_ = [g_*i*,1_,…, g_*i*,2*r*_] and g_*i,j*_ ∈ {0, 1} for any *i* and *j*. Let cn be the aggregate coverage vector observed from HTS data. We introduce a binary variable *z_i_* for each g_*i*_ that indicates if g_*i*_ is a part of the solution or not. We can model the objective—minimization of difference between the observed aggregate coverage and predicted solution—as follows:

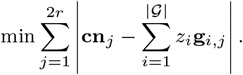

While this model performs well on WGS and targeted data (Numanagić *et al*., 2018), it is rather sensitive to deviations from the expected coverage distribution. It can also not properly handle the cases where the normalized aggregate coverage is not stable or uniform.^3^ Thus, Aldy 4 improves the original CNEP formulation by introducing additional optimization terms. This is done by modifying the original objective term and extending it with two additional terms, resulting in a three-term optimization objective.

The first term is the same as the original CNEP objective, but focuses only on the regions associated with the pharmacogene (and not its pseudogene):

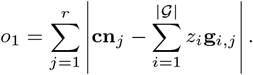

The second term of the objective function considers the interaction between the pharmacogene and the corresponding pseudogene region by considering the changes between their respective region coverage. For example, if the coverage of the exon 2 in *CYP2D6* is 3 and in *CYP2D7* is 2, the resulting region coverage difference would be 1. This difference can be further normalized (in this case, divided by 3). Using normalized differences allows us to handle samples in which the observed aggregate coverage (**cn**) varies between the regions due to various sequencing and alignment biases. Despite region-specific coverage variation, the relative abundances between the matching gene-pseudogene regions remain constant. This term can formally be expressed as:

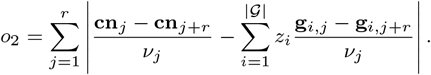

Here *ν_j_* = max{**cn**_*j*_, **cn**_*j*+*r*}_ + 1 is the normalization factor.

The final term of the objective function ensures that the ILP solver selects the most parsimonious solution:

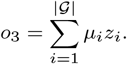

*μ_i_* is parsimony parameter (by default set to 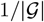). However, some unlikely configurations, such as left fusions, will have higher parsimony scores to reflect the observation that such configurations are rare (Sim *et al*., 2012).

Aldy 4’s modified CNEP model uses an ILP solver to minimize sum of these three terms *o*_1_ + *o*_2_ + *o*_3_. These solutions are passed to the later steps that will decide the best overall solution.

### 2.3 Star-allele calling

Aldy now proceeds by assigning the exact star-allele identifier to each of the *n* structural configurations obtained in the previous step. As stated in Section 2, a star-allele *S_i_* is defined as a tuple (g_*i*_, *A_i_*), where 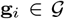 and 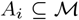. The star-allele assignment problem can also be modelled through the ILP as follows.

Let us indicate the presence of star-allele *S_i_* with a binary variable *a_i_*. Our goal is to select a set of star-alleles *S*_1_,…,*S_n_* such that (1) the set of the structural configurations that describes selected star-alleles is identical to the set the structural configurations from the previous step, and (2) the difference between predicted and observed coverage for each mutation *m* (denoted as cov(*m*)) is minimized. In other words, we want to minimize

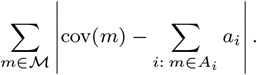

While conceptually simple, this model does not account for cases where database definitions are incomplete or incorrect. To account for these cases, we must allow the model to alter star-allele definitions if needed. Aldy thus introduces new binary variables *p_i,m_* and *q_j,m_* that indicate if a mutation m is to be “removed” from the star-allele *S_i_* (while being present in the database definition *A_i_*), or “added” to it (while being absent in *A_i_*). Then it attempts to minimize the following expression for each mutation *m*:

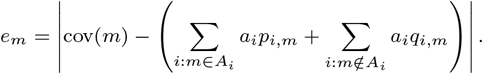

As *a_i_, p_i,m_* and *q_i,m_* are all binary variables, their product can be expressed as a set of linear constraints.

The minimization objective can be expressed as:

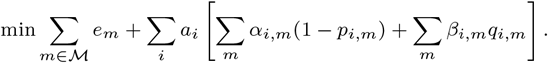

Parameters *α_i,m_* and *β_i,m_* are penalties for adding or removing the mutation *m* from allele *S_i_*. A novel mutation is less common than missing mutation, so generally we use *α_i,m_* = 2 and *β_i,m_* = 1 for any *i* and *m* (Numanagić *et al*., 2018). Note that not all mutations are the same: as functional mutations can fundamentally alter the behavior of a star-allele (and thus change its designation), we disallow removing functional mutations from any star-allele and allow adding novel functional mutations to the allele if and only if no other assignment is possible. This is done by setting the corresponding *α_i,m_* to a very large value.

The star-allele calling model also enforces other constraints: each functional mutation must be expressed by at least one allele, and each structural configuration must be expressed by at least one allele compatible with it. Finally, Aldy performs two rounds of star-allele calling for improved accuracy. In the first round, Aldy only considers functional mutations and identifies all star-allele combinations that explain the present functional mutations. Aldy then uses the second round to refine the calls from the first round by considering the silent mutations as well. It finally selects the star-allele with the best refinement score as the final call.

The formulation Aldy 4 uses for this step remains similar to the original model used in the older versions of Aldy. The single major difference is the change in the first (functional star-allele calling) round: Aldy 4 can now call star-alleles that contain novel functional mutations—a common case when gene databases are incomplete—if no other call can be made.

### 2.4 Read-based phasing

The above-described model essentially performs a variant of statistical phasing: it utilizes the database knowledge to select the most likely haplotypes that best explain the given observations from the data. While performing wellin practice (Numanagić *et al*., 2018), there are nevertheless cases when the aforementioned model produces multiple equally-likely calls. It is also unable to assign a novel mutation to a particular star-allele unambiguously. Finally, in sporadic cases, the above model can produce incorrect results. These challenges can be resolved with long reads that provide long-range phasing information. Aldy 4 newly incorporates the handling of long-range phasing information to the star-allele calling model as follows.

Suppose that we are given *z* fragments *R*_1_,…, *R_z_*, each fragment being defined by a set of mutations that it spans: 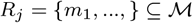. Each sequenced fragment originated from a single star-allele, and can thus be assigned to one of the star-alleles in the dataset. We can control this assignment by introducing a binary variable *f_i,j_* that is set if and only if a fragment *R_j_* is assigned to *S_i_*. Clearly, Σ_*i*_ *f_i,j_* must be one for every *R_j_* because each fragment originates from a single allele.

Ideally, we want to assign a *R_j_* to *S_i_* only if such an assignment agrees with the star-allele sequence as much as possible. In other words, we want to minimize the number of disagreements between allele *S_i_* and fragment *R_j_*. Thus, the total disagreement of an assignment can be expressed as follows:

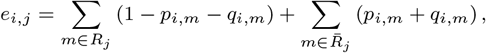

where 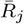 denotes the set of mutations that are not present in read *R_j_* but are spanned by it.

The total phasing error can be expressed as 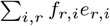. This expression can be added to the objective function of the star-allele calling model. Although the expanded version of this expression contains quadratic terms, each quadratic term is a product of two binary variables and, as such, can be trivially linearized.

As a final remark, note that the number of binary variables in the phasing model is dependent on the total number of present reads and alleles. In some cases, it can exceed half a million variables, making the overall model very costly to solve. The model can be significantly improved by using a smaller random sample of fragments, where the size of the random sample depends on the number of present reads and alleles.

### 2.5 Limitations

Aldy uses ILP solvers to solve the presented models. While ILP solving is NP-hard even when restricted to the models mentioned above (Numanagić *et al*., 2018), all these models are solvable in practice in less than a minute thanks to the state-of-the-art integer programming solvers utilized by Gurobi (Gurobi Optimization, LLC, 2022) or CBC (Forrest *et al*., 2018) solvers.

In some rare instances, Aldy cannot unambiguously call star-alleles from short-read datasets due to the read length limitations and lack of strand information. In these cases, Aldy will report all possible solutions. In some cases, this might be misleading; for example, a *68+*4/*5 call can be reported as *68/*4 (where *5 stands for deletion allele). However, both calls are functionally identical and should be treated as equal (as we do below). Aldy also makes heavy use of the existing star-allele databases to call star-alleles and fusion breakpoints. While it can handle cases where the database is incomplete or lacking, it can theoretically report incorrect results if a present allele is wildly divergent from any allele in the database.

Finally, Aldy 4’s detection of structural configurations is highly dependent on the stability of coverage across different sequencing runs. While this is not a significant issue for short-read WGS and targeted sequencing panels, the coverage might vary more than expected in PacBio samples. For this reason, Aldy 4 brings support for the exploration of a broader solution space when needed to account for potential noise.

## 3 Results

We have compared Aldy 4 (with PharmVar v5.1.15) against Astrolabe v0.8.7.2 (Twist *et al*., 2016), StellarPGx v1.2.5 (Twesigomwe *et al*., 2021), Stargazer v1.0.8 (Lee *et al*., 2019) and Cyrius v1.1.1 (Chen *et al*., 2021). We also compared Aldy 4 against the previous version of Aldy, Aldy v3.3 (Numanagić *et al*., 2018). The comparisons were done on a sizeable GeT-RM set of publicly available samples and genes for which genotyping panel validations were available (Pratt *et al*., 2010, 2016a; Gaedigk *et al*., 2019). These samples were sequenced with three technologies: (1) PGRNseq v.3 Illumina-based pharmacogene-targeted panel (Gordon *et al.* (2016); 137 samples), (2) Illumina whole-genome sequencing (WGS; 70 samples), and (3) 10X Genomics sequencing (95 samples). In addition to these samples, we also ran Aldy on the set of 45 Coriell samples sequenced by PacBio HiFi pharmacogene-targeted panel (Portik *et al*., 2021; Kingan *et al*., 2022) and validated by Scott *et al*. (2021).

Aldy 4 and other tools were run on the following 20 genes: *CYP1A1, CYP1A2, CYP2A13, CYP2A6, CYP2B6, CYP2C8, CYP2C9, CYP2C19, CYP2D6, CYP2F1, CYP2J2, CYP2S1, CYP3A4, CYP3A5, CYP3A7, CYP3A43*, *CYP4F2*, *DPYD*, *SLCO1B1*, and *TPMT*. While Aldy 4 also supports additional 15 genes, their evaluation was omitted because we did not have the ground truth panel data for these genes. Note that not every tool supports all of these genes: as a rule of thumb, Stargazer, Aldy 3 and Aldy 4 provide the broadest support, while the other tools are geared towards a small subset of these genes (typically CYP genes, such as *CYP2D6* and *CYP2C19*).

The available ground truth data is obtained through genotyping panels and assays designed to detect only the common *major star-alleles* (i.e., alleles defined solely by functional variants). These panels often cannot call minor star-alleles (i.e., alleles defined by non-functional variants and functionally indistinguishable from the major star-alleles), as well as less common alleles. The low resolution of the available ground-truth data and the differences in database specifications between the different tools necessitated a few accommodations within the evaluation process for the sake of fairness. First, we updated ground truth calls that missed the presence of less common variants and alleles. Updates were only done if there was a consensus between the star-allele calling tools that differed from the ground truth data and if an updated call extended the validated allele definition (i.e., if the mutations defining the validated allele also form a part of the consensus definition). Note that a similar approach was used in Numanagić *et al*. (2018). Each updated call was further manually inspected to ensure that the variants missing from the ground truth calls are indeed present and not sequencing artifacts. In rare instances, it was hard to precisely distinguish the presence of the variant, especially if the variant allele frequency (VAF) was too low (alleles with lower VAFs are sometimes caused by the sequencing or read alignment bias, especially in the presence of pseudogenes, and are typically validated through Sanger sequencing). Samples with such variants were marked as “Need Validation” (Table 1). For such samples, calls that either used or ignored such ambiguous variants were deemed “correct”. Second, we have followed the common strategy employed in clinical studies (Ly *et al*., 2022) by only comparing the major star-allele calls and ignoring the minor star-allele designations. In other words, only the phasing of functional variants was considered; non-functional and silent variants that do not alter the functionality of an allele were ignored (i.e., a *1A/*2B minor star-allele call was treated as a functionally equivalent *1/*2 major star-allele call^4^).

**Table 1.**
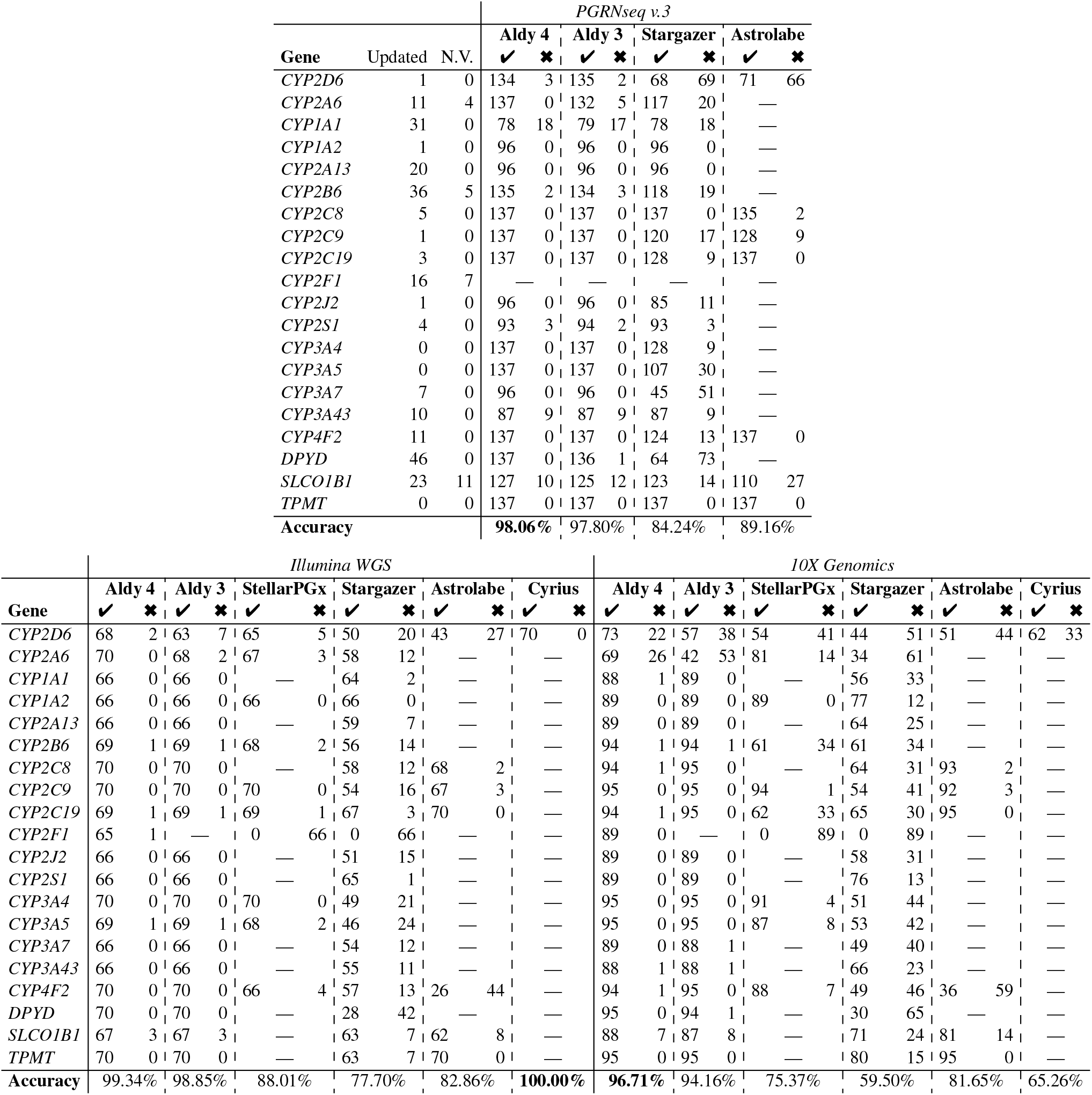
Summary of the star-allele calls generated by six tools (Aldy 4, Aldy 3, StellarPGx, Stargazer, Astrolabe and Cyrius) on 137 GeT-RM samples sequenced by three different technologies. Bold results indicate the best tool for a given gene. Some tools do not support all genes; those cases are indicated with a dash (**—**). Checkmark (**✓**) indicates the call that matches the updated validation panel star-allele call; a cross-mark (**✖**) indicates the mismatch. Number of updated panel calls, as well as the calls that need further validation (marked with N.V.), is indicated at the beginning. Note that the total number of samples varies across genes and technologies due to the availability of sequencing data and ground truth validation. Detailed results are available at https://github.com/0xTCG/aldy.

Where possible, we used the *CYP2D8* region as the copy number neutral region; exceptions include Aldy 4 using *F1* region for the PacBio data. Some tools, such as Astrolabe and Stargazer, relied on VCF files; where needed, VCFs were generated by `bcftools` (Li *et al*., 2009).

All results were obtained on machines with Intel Xeon E5-2680v4 and 8260 CPUs. Each evaluated tool genotypes a single gene in a single sample within a few minutes, regardless of the sequencing technology used. However, note that Aldy 4 only needs BAM/CRAM to run; other tools often require VCF or GDF files that can take significant time to generate.

Overall, the best accuracy on short-read datasets (PGRNseq v.3, Illumina WGS and 10X Genomics) was achieved by Aldy 4 (98.04%), followed by Aldy 3 (96.93%), Astrolabe (84.56%), Cyrius (82.63%), StellarPGx (81.69%) and Stargazer (73.82%).

### 3.1 PGRNseqv.3

Aldy 4, Aldy 3, Stargazer and Astrolabe were run on 137 PGRNseq v.3 targeted sequencing (Gordon *et al*., 2016) samples from the GeT-RM collection. PGRNseq v.3 targets common pharmacogenes and sequences them at high depths (up to 1000 × per loci).

Note that we could not get either Stargazer or Astrolabe to run on targeted sequencing data natively; for that reason, we used VCF files as an input for these tools. Because of the limited nature of VCF files, these tools were unable to call copy number changes and fusions on this dataset.^5^ We have omitted the comparison with StellarPGx as it does not support targeted sequencing data.

As can be seen in Table 1, Aldy 4 identifies nearly all of the alleles in all genes correctly—more than the other two tools—with a total accuracy of 96.7%. In some cases (e.g., failed cases in the genes *CYP1A1* and *CYP2B6*), no caller was able to call correct star-alleles because the PGRNseq panel did not sequence the variant of interest (e.g., a non-exonic downstream variant rs4646903 that defines *CYP1A1*2A* was not covered by the panel at all).

On this dataset, Aldy 4’s performance is only marginally better than Aldy 3. This is expected as neither of the model updates unique to Aldy 4 applies to the high-quality PGRNseq dataset with stable coverage. Minor changes are mostly due to the differences in the variant calling (e.g., Aldy 4’s incorporation of quality scores and mapping qualities).

### 3.2 Illumina WGS

We have run all tools on 70 Illumina HiSeq-sequenced WGS samples from the GeT-RM sample collection. These samples were sequenced with an average depth of roughly 30×. The details are also available in Table 1.

Here, Aldy 4 again calls nearly all star-alleles correctly and genotypes more samples than the competition for every considered gene. The only exception is *CYP2D6,* where Cyrius genotypes one sample (NA21781) more than Aldy 4. In this case, Aldy 4 identifies the *2 allele as *65; however, the *65 allele extends the *2 allele with a single SNP (rs1065852), and it is unclear if this allele is indeed a *2 or a *65.

Aldy 4 and other tools were able to correctly call alleles defined by intronic and downstream variants across the genes on this data. Note that the main reason behind the Stargazer’s lower accuracy on this dataset was copy number calling: while Stargazer often identified the star-allele correctly, it would often call them more times than needed (e.g., *1/*2+*2 instead of *1/*2).^6^

Note that Astrolabe used a modified *CYP4F2* database whose allele nomenclature differed from the other databases. Thus, we have omitted comparison with Astrolabe on *CYP4F2* for the sake of consistency. We also observed a large number of mismatches in *SLCO1B1* across all tools due to the incomplete panel validation and inconsistent database specifications used by various tools.

Finally, the improvements in the copy number model and more sensitive variant calling in Aldy 4 account for a few improved calls on more complex *CYP2D6* and *CYP2A6* samples.

### 3.3 10X Genomics

We have run all tools on 95 GeT-RM samples sequenced by a 10X Genomics WGS sequencer. The average depth of sequencing was roughly 40×. Because several important pharmacogenes reside within repeated regions of the human genome, we used EMA aligner (Shajii *et al*., 2018) in density-based optimization mode for improved alignment of the 10X reads to the reference genome (hg19). The comparison details are available in Table 1.

Although the 10X Genomics protocol uses Illumina HiSeq for sequencing, the read coverage is not as uniform as it is in an average Illumina WGS sample. 10X-specific biases also result in quite a few misaligned reads compared to the WGS data. For this reason, the overall allele calling accuracy is lower than the WGS dataset; this is especially evident in Stargazer, where the accuracy of its copy number detection module is even lower than in WGS data.

However, Aldy 4 still correctly calls the majority of alleles (with 95.9%) accuracy, especially when compared to the other tools. The most challenging genes for all tools were *CYP2A6* and *CYP2D6.* Aldy’s accuracy is lower in these genes, primarily due to the occasional copy number mismatch (due to the coverage unevenness) and sequencing artifacts (where many mis-identified variants were either an artifact or were under-sequenced). Note that Aldy 4 benefited from the novel phasing module that was able to successfully utilize 10X Genomics barcodes to link long-distance variants together. Finally, we observe the significant improvements over Aldy 3 in *CYP2D6* and *CYP2A6* samples on this dataset due to an improved copy-number model that better handles noisy coverage and ambiguous variants (a common case in 10X Genomics samples), and is as such able to improve the calling accuracy up to 30% in these genes.

### 3.4 PacBio HiFi

Finally, we ran Aldy 4 on two sets of PacBio HiFi samples sequenced by a custom targeted pharmacogenomics panel (Portik *et al*., 2021; Kingan *et al*., 2022). The first set contained 24 samples, while the second set was comprised of 21 samples. The region gene coverage of these datasets varies—it can be as low as 10 ×—and at times exceed even 200 ×. We compared Aldy’s calls with those of Astrolabe. While none of the other tools support PacBio long reads natively, we were able to at least run Astrolabe in VCF mode. The validation data was obtained from Scott *et al.* (2021) and Pratt *et al*. (2016b). The call details are available in Table 2.

**Table 2.**
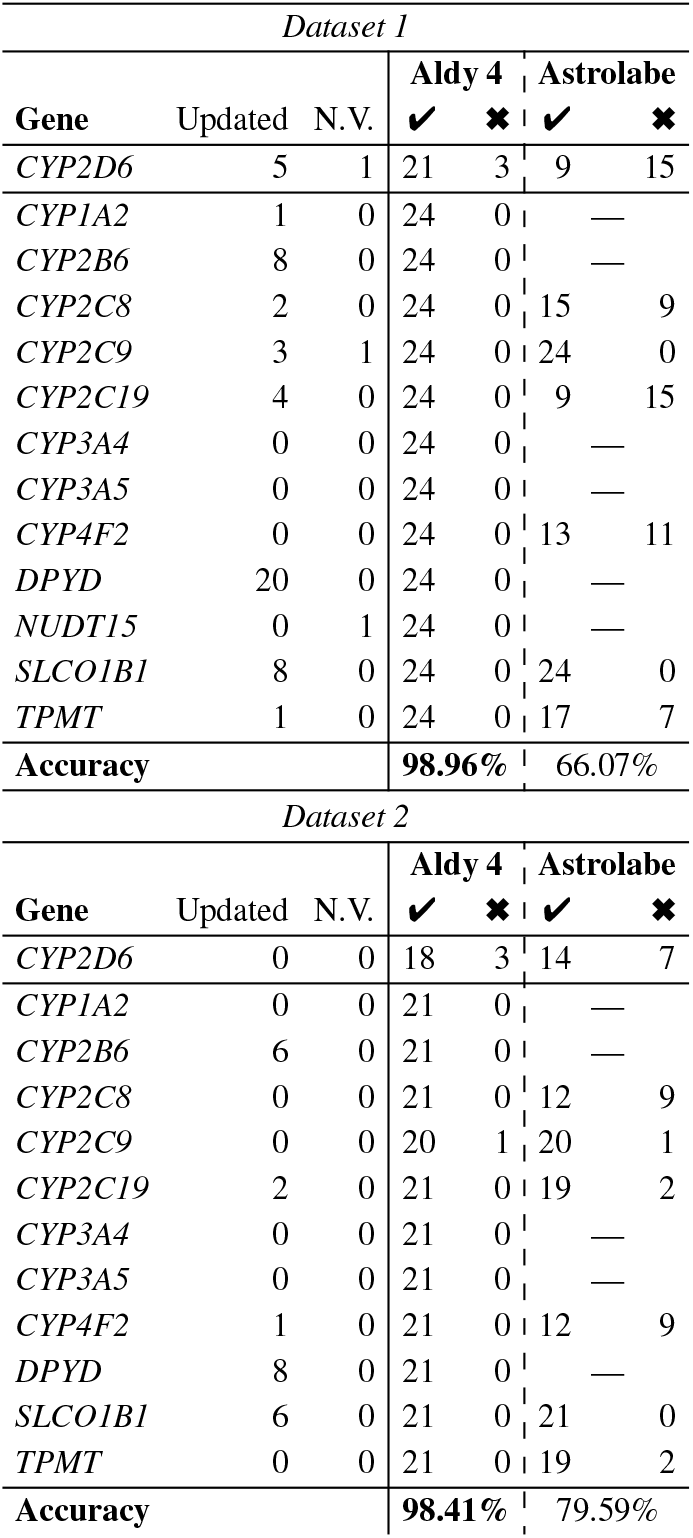
Overview of the star-allele calls generated by Aldy 4 and Astrolabe on PacBio HiFi targeted data. Some tools do not support all genes;those cases are indicated with a dash (—). Check mark (**✓**) indicates the call that matches the updated validation panel star-allele call;a cross-mark (**✖**) indicates the mismatch. Number of updated panel calls, as well as the calls that need further validation (marked with N.V.), is indicated at the beginning.

Star-allele calls generated by Aldy 4 agree with the ground truth in all genes except for a few *CYP2D6* calls and one *CYP2C9* call. Furthermore, its calls augmented and phased many ground-truth calls generated by panels with limited SNP coverage with additional SNPs observed by PacBio data (Table 2). Aldy was also able to find and phase novel alleles that have not been cataloged in genes *CYP2B6, CYP2C19, CYP3A4, DPYD* and *SLCO1B1.* Further validation of such novel calls, as well as of the calls that were deemed ambiguous, is needed to fully confirm and understand such alleles.

When it comes to *CYP2D6,* Aldy 4’s calls disagree with the groundtruth data due to the difference in predicted copy number. In two instances, Aldy 4 called an additional copy (i.e., *1+*1 instead of *1, and *4+*4 instead of *4), while in the other two instances, Aldy 4 did not call an existing copy (i.e., it called *2 instead of *2+*2 and *10 instead of *10+*10). In one instance, Aldy called *36 instead of *10 (note that these alleles are nearly identical); in the final instance Aldy did not call nonfunctional *68 fusion allele. In all these cases, the observed coverage was noisy, and further validation is needed to ascertain the exact copy number of these samples. We would also like to point out that Astrolabe’s calls in genes *CYP2C19* and *SLCO1B1*, as well as on *CYP2D6* on the second dataset, were highly ambiguous, often containing more than ten functionally different solutions.

### 3.5 Other remarks

We have observed that many tools often confuse *CYP2B6*6* and *CYP2B6*9* alleles that differ only in rs2279343 SNP. This SNP is often either under-sequenced or covered by ambiguous reads that potentially originate from the neighboring *CYP2B7* pseudogene, and is thus hard to call with high confidence in some technologies (e.g., PGRNseq v.3). When the true call was ambiguous, we have deemed both possible calls “correct”. Similar cases were also observed with *CYP2A6*1* and *CYP2A6*35* alleles. Further validation is needed to properly ascertain the true existence of these alleles in problematic samples.

If multiple allele calls were generated by a tool for a given sample and gene combination, the call was deemed “correct” if at least one such multi-call matched the ground truth. Note that the prevalence of multiple calls was overall low: around 1.2% for Aldy 4, 2.7% for Aldy 3, 1.8% for Stargazer, 1.8% for StellarPGx and 15.7% for Astrolabe. Aldy 4’s new phasing model was a significant factor for a low multi-call rate: while the rate was 1.8% on PGRNseq v.3 and 1.9% on WGS samples due to the short read lengths of such samples, it decreased to 0.5% on 10X and PacBio samples that allowed better phasing. The vast majority of ambiguous calls were observed when genotyping *CYP4F2* and *SLCO1B1*.

## 4 Conclusion

Pharmacogenomics is becoming a key component of evidence-based medicine (Relling and Evans, 2015). Genes like *CYP2D6* and *CYP2C19* regulate a large portion of clinically prescribed drugs; other genes, such as *HLA* or *IGH* gene cluster, are vital for understanding the immune response (Ford *et al*., 2020, 2022). As their function is dependent on their haplotype, it is of vital importance to genotype and haplotype these genes prior to administering medical treatment (Crews *et al*., 2014). High-throughput sequencing technologies are a natural candidate for this process, especially when considered that the currently available clinical genotyping panels are often restricted only to the most common genotypes and struggle to detect more complex structural altercations within pharmacogenes.

In this work, we have presented Aldy 4, the first tool that can accurately and consistently call star-alleles in data from various sequencing technologies, including but not limited to long-read PacBio data, Illumina short-read sequencing in all of its flavors (i.e., wholegenome, whole-exome, and targeted capture data), as well as the 10X Genomics barcoded data. Aldy 4 achieves this by employing combinatorial optimization models to solve various challenges associated with calling pharmacogenetic haplotypes from sequencing data, such as copy number and structural variation detection, variant calling and variant phasing, ultimately resulting in a star-allele decomposition of a gene of interest. We have shown the strength of Aldy 4’s approach through a series of comparisons against the current state-of-the-art star-allele callers, in which Aldy 4 always performed the best. We hope that Aldy 4 will be of vital importance to clinicians in tailoring prescription recommendations, thus ultimately leading to improved medical care.

There are still some open questions left that need to be answered in future work. Most importantly, the panel-validated calls improved by the star-allele callers through the use of HTS data—often containing novel alleles not previously cataloged in the existing databases—need to be validated in a wet-lab environment for all genes presented, as was done recently for a selection of CYP2C genes (Gaedigk *et al*., 2022). More tests are also needed on larger cohorts to accurately evaluate the precision of these tools, Aldy 4 included, on rare fusions. The incorporation of other highly polymorphic pharmacogenomics regions, such as *HLA* or *IGH*, should also be considered, as Aldy (and other evaluated pharmacogenomics tools) are currently unable to handle the complexities of such regions. Finally, the complete characterization of minor star-alleles, accompanied by the careful characterization of non-coding variants, is also needed to understand the full effect of pharmacogenes on the treatment and drug dosage decisions.

## Acknowledgements

We would like to acknowledge Steven E. Scherer and Xiang Qin from Human Genome Sequencing Center at Baylor College of Medicine (Houston, Texas) for providing us with PGRNseq v.3 samples.

## Footnotes

1 https://github.com/PacificBiosciences/pbmm2

2 For the sake of clarity, here we present an idealized version where each structural configuration can be selected only once. In practice, Aldy allows selecting the same configurations multiple times.

3 For targeted panels with non-uniform coverage distributions, aggregate coverage can be “normalized” by dividing it by the coverage of the control sample if it is stable across different samples. Aldy does this automatically for known targeted panels.

4 Major star-alleles are typically distinguished by the number (e.g., *1 functionally differs from *2). Minor star-alleles are typically distinguished by a letter (e.g., *2A and *2B harbor different silent variants despite sharing common functional variants).

5 While Stargazer has a mode for targeted data, we were unable to get good results with it.

6 Note that Aldy only calls copy numbers and fusions on genes that are known to harbor such changes; otherwise, it assumes that two copies are present.

